# Regulatory Genomics of Preeclampsia-Specific Risk Variants Highlights Immune and Endothelial Mechanisms

**DOI:** 10.64898/2026.05.26.728031

**Authors:** Mohammed A. Farahat, Malak Abbas, Gideon A. Wiafe, Taneisha Gillyard Cheairs, Melissa Nel, Amadou Gaye

**Affiliations:** Department of Integrative Genomics and Epidemiology, School of Graduate Studies, Meharry Medical College, Nashville, TN, USA; National Institute of Diabetes and Digestive and Kidney Diseases, National Institutes of Health, Bethesda, MD, USA; Clinical Omics and Informatics Unit, Department of Medicine, Neuroscience Institute, University of Cape Town, South Africa; Department of Biomedical Sciences, School of Graduate Studies, Meharry Medical College, Nashville, Tennessee, USA; Department of Computer Science, Higher Future Institute for Specialized Technological Studies, Cairo, Egypt; African Bioinformatics Institute

## Abstract

**Background:** Preeclampsia (PE) is a complex hypertensive disorder of pregnancy characterized by endothelial dysfunction, immune dysregulation, and systemic vascular injury. Multiple genome-wide association studies (GWAS) have revealed genetic signals shared with hypertension and blood pressure traits, potentially obscuring biological mechanisms that are more specific to PE pathogenesis. Furthermore, the functional consequences of most PE-associated variants remain poorly understood. In addition, GWAS relies on short-read sequencing and array-based analyses, limiting the ability to identify insertions, deletions, and other structural variants that may contribute to disease-associated regulatory mechanisms. In this study, we investigated the regulatory architecture of PE-specific genetic variants and evaluated their potential linkage disequilibrium (LD) with structural variants.

**Methods:** We integrated GWAS, transcriptomic, and long-read sequencing data to investigate the regulatory architecture of PE-specific genetic variants. Summary statistics for PE, hypertension, systolic and diastolic blood pressure were obtained from the GWAS Catalog, and variants uniquely associated with PE (P ≤ 1×10^-4^) were prioritized. Cis-expression quantitative trait locus (cis-eQTL) analyses were performed in whole-blood RNA-sequencing data from 180 African American women. Significant associations were replicated in biologically relevant tissues from the GTEx Project, including vascular, renal, and immune-related tissues. Long-read sequencing-derived structural variants (SVs) were subsequently evaluated for LD with replicated eQTL loci.

**Results:** A total of 10,843 PE-specific variants, present in whole-genome sequencing data of the 180 women, were evaluated. Cis-eQTL analyses identified 480 significant eQTL-gene associations involving 277 unique variants and 192 genes (FDR ≤ 0.05). Replication analyses supported 69 eQTL-gene associations across five GTEx tissues, involving 35 variants and 14 genes. Replicated signals were enriched in vascular tissues, particularly artery tibial and artery aorta. Several prioritized genes converged on immune and vascular pathways, including MICA, HLA-DPB1, SEMA4D, JUP, ZFP57, and TMEM204. Integration of GWAS and eQTL effects demonstrated consistent regulatory shifts associated with PE-risk alleles, including downregulation of immune-related loci and upregulation of select vascular-associated genes. Long-read sequencing analyses identified 66 high-LD (r^2^ ≥ 0.80) SNP-SV-gene associations, including 12 replicated eQTL variants, 8 candidate SVs, and 3 replicated genes, suggesting that structurally complex genomic regions may contribute to the observed regulatory signals.

**Conclusions:** The tissues enriched in the regulatory signal highlight the importance of systemic endothelial biology in PE susceptibility. The findings of this study support a model in which PE-specific genetic susceptibility converges predominantly on interconnected immune and vascular regulatory mechanisms. The integration of eQTL analyses with long-read structural variant discovery provides additional insight into the complex genomic architecture underlying PE and highlights candidate regulatory loci that may not be adequately captured through conventional GWAS approaches alone. The study also emphasizes the importance of conducting functional genomic analyses in diverse populations to improve understanding of disease biology and advance precision medicine efforts.

**GRAPHICAL ABSTRACT:** 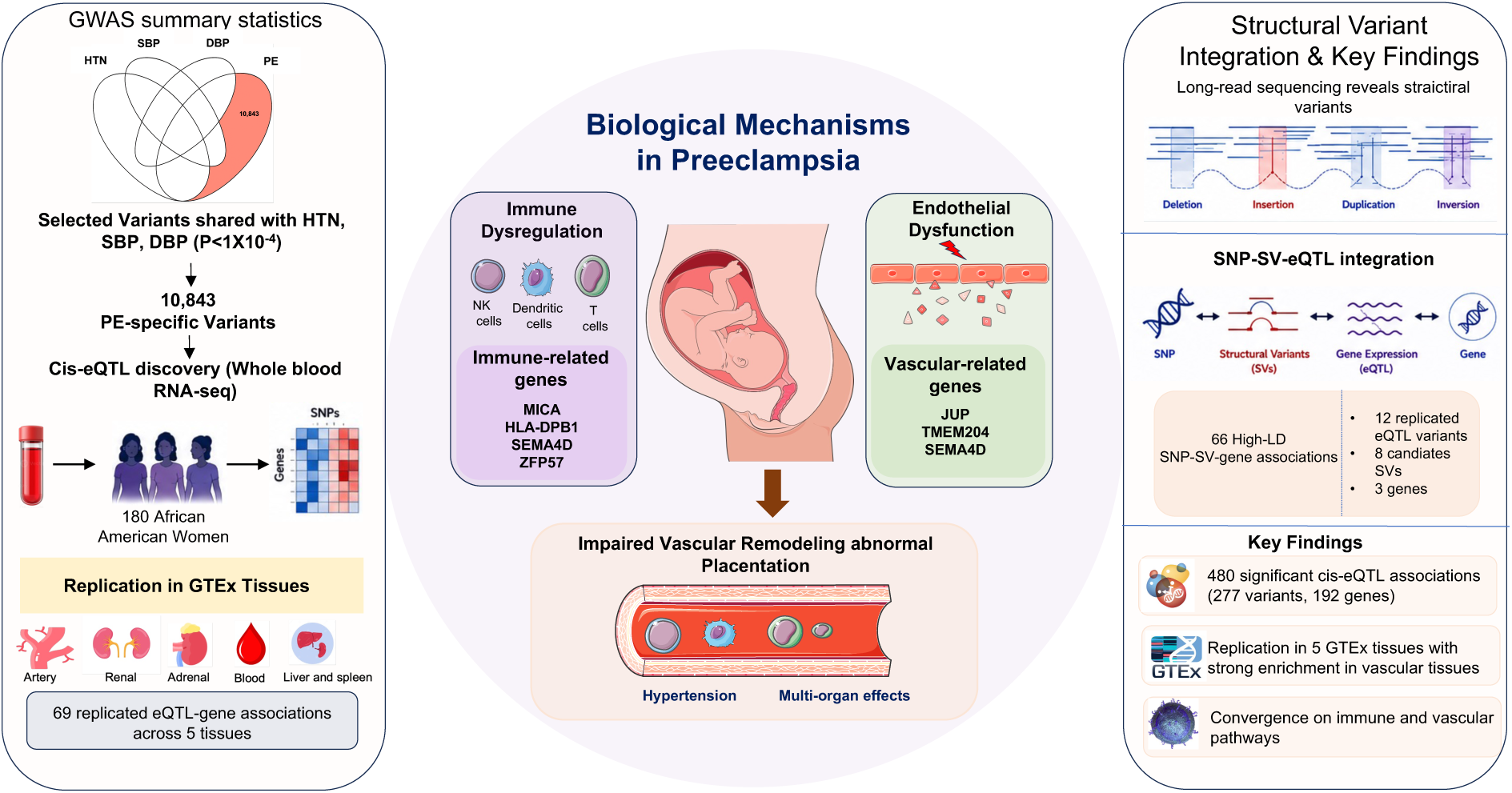

Regulatory Genomics of Preeclampsia-Specific Risk Variants Highlights Immune and Endothelial Mechanisms.

GWAS summary statistics for preeclampsia, hypertension, SBP, and DBP were integrated to identify 10,843 preeclampsia-specific variants that were subsequently evaluated in cis-eQTL analyses using whole-blood RNA-sequencing data from 180 African American women (left). Cis-eQTL analyses identified 480 significant associations involving 277 variants and 192 genes (FDR ≤ 0.05), of which 69 eQTL-gene associations involving 35 variants and 14 genes replicated across five GTEx tissues, with strongest enrichment observed in vascular tissues, particularly artery tibial and artery aorta (center). Prioritized genes, including MICA, HLA-DPB1, SEMA4D, JUP, ZFP57, and TMEM204, converged on interconnected immune and endothelial pathways associated with systemic vascular dysfunction, impaired placentation, and inflammatory dysregulation in preeclampsia. Integration of long-read sequencing data further identified 66 high-LD SNP-SV-gene associations involving 12 replicated eQTL variants, 8 candidate structural variants, and 3 replicated genes, suggesting that structurally complex genomic regions may contribute to regulatory mechanisms not fully captured through conventional GWAS approaches alone.

eQTL indicates expression quantitative trait locus; FDR, false discovery rate; GTEx, Genotype-Tissue Expression project; SBP, systolic blood pressure; DBP, diastolic blood pressure; LD, linkage disequilibrium; SV, structural variant.

## INTRODUCTION

Preeclampsia (PE) is a complex multisystem disorder of pregnancy characterized by new-onset hypertension and evidence of maternal organ dysfunction, typically arising after 20 weeks of gestation ^1, 2^. Affecting approximately 2-8% of pregnancies worldwide, PE remains a major contributor to maternal and fetal morbidity and mortality, particularly in low-resource settings and among women of African ancestry ^3, 4^. Beyond its immediate obstetric consequences, PE is increasingly recognized as a long-term cardiovascular and metabolic risk factor for both mothers and offspring ^5–7^. Despite decades of research, the molecular mechanisms underlying PE remain incompletely understood, limiting the development of effective biomarkers and targeted interventions ^8–10^.

The pathophysiology of PE is multifactorial and involves abnormal placentation, endothelial dysfunction, immune dysregulation, oxidative stress, and systemic vascular injury ^11, 12^. These biological processes overlap substantially with mechanisms implicated in hypertension and blood pressure regulation, supporting the hypothesis that PE shares part of its genetic architecture with cardiometabolic and vascular traits ^10, 13^. Genome-wide association studies (GWAS) have identified multiple loci associated with PE susceptibility, including variants mapping to immune-related, angiogenic, and vascular regulatory pathways ^14–16^. However, most GWAS signals reside in non-coding regions of the genome, making it difficult to determine their functional consequences and causal target genes ^17, 18^.

Expression quantitative trait locus (eQTL) analysis provides a powerful framework for linking disease-associated variants to downstream transcriptional regulation ^19, 20^. By integrating GWAS findings with transcriptomic data, eQTL analyses can help identify regulatory variants that influence disease susceptibility through altered gene expression ^21–23^. This strategy has proven particularly informative for complex cardiovascular and immune-mediated disorders, where regulatory variation plays a central role in disease biology ^24, 25^. In PE, however, functional genomic studies remain relatively limited, especially in populations of African ancestry, which continue to be underrepresented in large-scale genomic research despite experiencing disproportionately high disease burden ^26–28^.

Most prior studies investigating the genetics of PE have focused primarily on variants shared with hypertension or blood pressure traits, reflecting the strong clinical overlap between these conditions ^14, 29^. While this approach has yielded important insights into shared vascular biology, it may obscure regulatory mechanisms that are more specific to PE pathogenesis. Identifying variants uniquely associated with PE may therefore help disentangle disease-specific biological pathways from broader blood pressure-related processes. Furthermore, understanding how these variants regulate gene expression across vascular, immune, and renal tissues may provide important insights into the maternal systems most strongly involved in disease development.

An additional limitation of conventional GWAS and short-read sequencing approaches is their reduced ability to capture structural variants (SVs), including insertions and deletions, which represent a substantial and functionally important component of human genetic variation ^30, 31^. Structural variants can alter gene regulation through multiple mechanisms, including disruption of promoters, enhancers, chromatin organization, and transcription factor binding sites ^31, 32^. Emerging evidence suggests that SVs contribute significantly to regulatory variation and may underlie disease-associated signals detected by nearby single nucleotide polymorphisms (SNPs) through linkage disequilibrium ^31, 33^. However, the contribution of SVs to PE-associated regulatory loci remains largely unexplored, particularly in African ancestry populations where genomic diversity is greatest and structural variation is comparatively understudied ^34^.

Recent advances in long-read sequencing technologies now enable improved detection and characterization of SVs compared with conventional short-read platforms. Integrating long-read SV discovery with SNP-based eQTL analyses provides an opportunity to refine the interpretation of GWAS loci and identify candidate causal variants underlying observed regulatory associations. Such approaches may be especially valuable for uncovering complex regulatory architectures in regions enriched for immune and vascular regulatory genes, including the major histocompatibility complex (MHC), which has repeatedly been implicated in PE susceptibility ^35, 36^.

In this study, we integrated GWAS, transcriptomic, and long-read sequencing data to investigate the regulatory architecture of PE-specific genetic variants. First, we identified variants associated uniquely with PE, excluding those shared with hypertension and blood pressure traits. We then performed cis-eQTL analyses in whole blood transcriptomic data from African American women to identify regulatory relationships between PE-specific variants and nearby genes. Significant findings were subsequently evaluated in biologically relevant tissues from the GTEx Project, including vascular, renal, and immune-related tissues^37^. Finally, we leveraged long-read sequencing-derived structural variant data to assess whether replicated eQTL loci were linked to candidate SVs in strong linkage disequilibrium. Together, these analyses provide a multi-layered framework for identifying regulatory mechanisms underlying PE susceptibility and highlight immune and vascular pathways that may contribute to disease pathogenesis.

## MATERIALS AND METHODS

### Study Cohorts

#### - GENE-FORECAST

The Genomics, Environmental Factors and the Social Determinants of Cardiovascular Disease in African Americans Study (GENE–FORECAST) is a population–based research initiative aimed at generating comprehensive phenotypic data from African American adults using an integrated multi–omics approach. The study population consists of U.S.-born African American individuals between 21 and 65 years of age, recruited primarily from communities in the greater Washington, DC, metropolitan area through a community–engaged recruitment framework, as described previously ^38^. The research protocol received approval from the Institutional Review Boards at both the National Institutes of Health and Meharry Medical College. All participants provided written informed consent prior to enrollment, and all study activities were conducted in compliance with applicable institutional policies and regulatory standards.

#### - NeuroSeq

The NeuroSeq study is a genomic research study led by the Clinical Omics and Informatics Unit at the University of Cape Town’s Neuroscience Institute that recruits individuals of diverse ancestries presenting with neurological phenotypes of both presumed genetic and environmentally influenced origin across participating clinical sites in Africa. While recruitment is ongoing, the current sample comprises predominantly African-ancestry adult individuals with amyotrophic lateral sclerosis recruited through the ALSAfrica clinical site network. At the individual level, the NeuroSeq study is observational and primarily focused on describing genetic and epigenetic findings in individuals with neurological phenotypes. As recruitment expands, the accumulated data will enable case-control and cohort-based comparisons within the study population and against external datasets. Structural variants were ascertained from long-read HiFi sequencing data generated from this continental African cohort. Although the cohort was recruited for an unrelated phenotype (ALS), linkage disequilibrium is a population-level property determined by ancestral recombination and haplotype structure and is estimated independently of disease status. Therefore, the suitability of the cohort as a haplotype reference depends primarily on ancestral representativeness rather than phenotypic ascertainment. Furthermore, the loci evaluated in this study are not established ALS-associated loci and no known genetic overlap between ALS and preeclampsia has been reported at these regions, so phenotypic ascertainment is not expected to bias LD at the loci of interest. The NeuroSeq study was approved by the University of Cape Town’s Faculty of Health Sciences Research Ethics Committee (HREC REF 920/2025). All participants provided written informed consent to participate, including the use of aggregate, de-identified data in non-disease-specific studies as control data.

### Genotype Data

#### - Short-read sequencing data

Genomic DNA was extracted from peripheral blood samples collected in EDTA–treated tubes and quantified using the PicoGreen fluorescence assay. Whole–genome sequencing libraries were prepared using a PCR–free protocol with small insert sizes. Sequencing was carried out on the Illumina NovaSeq 6000 platform, generating 151–base–pair paired–end reads with a mean coverage depth of approximately 30x. Raw sequencing data were processed using the Broad Institute’s Whole Genome Germline Variant Discovery workflow. Reads were aligned to the human reference genome build 38 with the Burrows-Wheeler Aligner. Variant discovery, gVCF generation and aggregation, joint genotyping, and downstream quality control were performed in accordance with GATK4 best–practice guidelines. Among participants with available whole–genome sequencing data, a subset of 180 women also had corresponding whole–blood transcriptomic profiles obtained through mRNA sequencing.

#### - Long-read sequencing data

A subset of the NeuroSeq cohort was analyzed in this study, comprising 30 individuals who underwent long-read sequencing using the PacBio Revio platform at Inqaba Biotec (Pretoria, South Africa). The cohort was predominantly composed of South African Black participants (SAB; n = 22), with additional samples from Nigeria (n = 5), Kenya (n = 2), and Zimbabwe (n = 1). Each sample was sequenced to an average target coverage of 30×. Across the cohort, the mean read length was 12.1 kb, with an average read quality score of 37.1. Sequencing output per sample averaged approximately 7.29 million reads, corresponding to a total yield of ∼92.2 Gb.

Structural variant (SV) discovery and genotyping were performed using a two-stage pipeline implemented in the PacBio structural variant caller pbsv (v2.10.0), optimized for long-read data. In the first stage, SV signatures were identified independently for each sample using the pbsv discover module with HiFi-optimized parameters applied to BAM files aligned to the GRCh38 reference genome. To improve the representation of variants in repetitive regions, a tandem repeat finder (TRF) BED file was incorporated during this step to normalize SV signatures. This process generated compressed SV signature files for each sample. In the second stage, these per-sample signatures were aggregated and jointly analyzed using pbsv call, enabling cohort-wide SV discovery and genotyping against the GRCh38 reference genome.

### Transcriptomics Data

GENE-FORECAST whole–blood gene expression profiles were obtained through messenger RNA sequencing (mRNA–seq). Total RNA was extracted from blood samples preserved in stabilization tubes using the MagMAX™ RNA isolation kit (Life Technologies, Carlsbad, CA). Library preparation was performed from total RNA using Illumina TruSeq reagents, producing indexed complementary DNA (cDNA) libraries following ribosomal RNA depletion.

Paired–end RNA sequencing was conducted using Illumina HiSeq 2500 and HiSeq 4000 platforms, with each sample sequenced to a depth of at least 50 million reads. Gene expression levels were quantified using the Broad Institute’s GTEx RNA–seq processing workflow, following their publicly available analysis pipeline ^39^. Transcripts exhibiting low expression were filtered out prior to normalization, defined as those with fewer than 2 counts per million (CPM) detected in at least three samples. Normalization of expression data was performed using the Trimmed Mean of M–values (TMM) method, which is appropriate for count–based RNA–sequencing data ^40^. Principal component analysis was subsequently applied to identify expression outliers, resulting in the removal of four transcripts from downstream analyses.

Following quality–control procedures, a total of 17,947 protein–coding genes were retained for subsequent analyses. Expression quantitative trait locus (eQTL) analyses were conducted in a subset of 180 women with both whole–genome sequencing and mRNA–sequencing data available.

### eQTL Analysis

The analytical workflow comprised three major components (**Figure 1**). We first retrieved summary statistics from large-scale GWAS of hypertension, SBP, DBP, and preeclampsia available in the GWAS Catalog. Genetic variants associated with these traits at p ≤ 1 × 10⁻⁴ were extracted, and variants associated only with preeclampsia were identified for downstream analyses. For all prioritized variants, we evaluated their potential regulatory effects on nearby genes within a 1 Mb window (cis region) using eQTL analyses.

**Figure 1:**
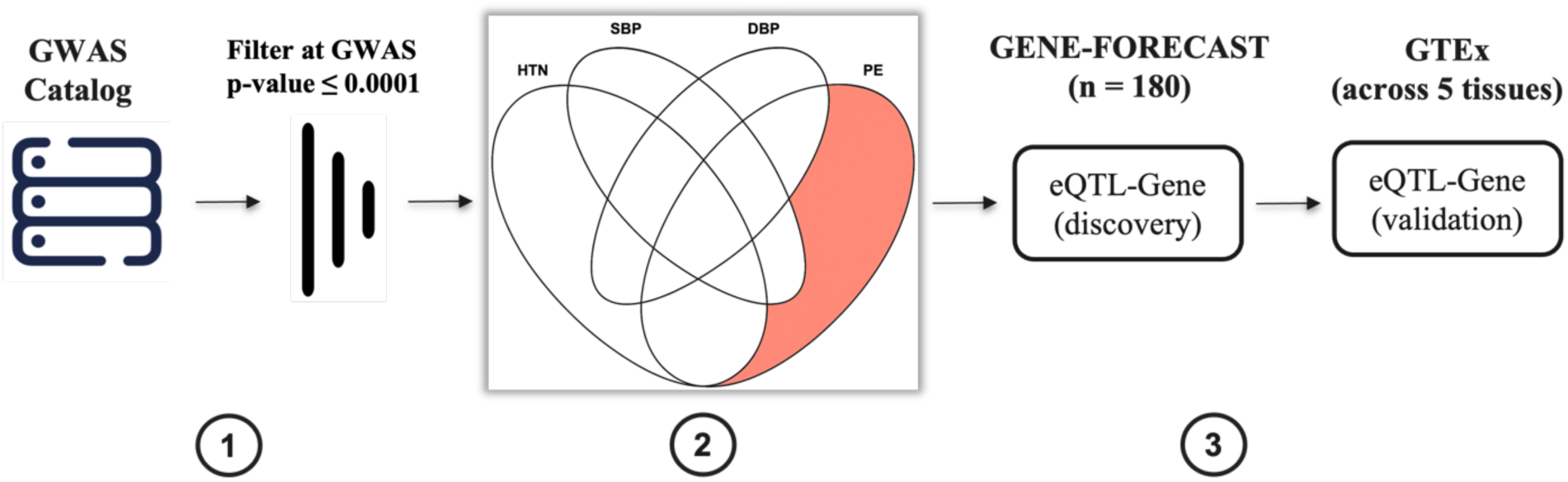
Study design and analytical workflow for identifying preeclampsia-specific regulatory variants and target genes. (1) Summary statistics were obtained from the GWAS Catalog and filtered to retain associations with P ≤ 1×10⁻⁴, followed by cross-trait comparison (HTN, SBP, DBP, and PE) to isolate variants uniquely associated with preeclampsia (PE-specific GWAS signals). (2) These PE-specific variants were integrated with whole-blood mRNA expression data in 180 women (GENE-FORECAST cohort) to perform cis-eQTL analyses and identify variant–gene regulatory relationships. (3) Significant eQTL-gene associations were subsequently validated in independent datasets from GTEx across whole blood and four additional relevant tissues to assess cross-tissue reproducibility.

Cis-eQTL analyses were conducted in whole blood using RNA-seq data of 180 women, from the GENE-FORECAST cohort, to determine whether shared GWAS variants influence gene expression. This approach provides insight into the regulatory mechanisms by which genetic loci may contribute to the pathophysiology of preeclampsia.

Analyses were performed using the R package *MatrixEQTL*, fitting models with normalized mRNA expression as the dependent variable and genotype dosages as the independent variable. Models were adjusted for age, and the top six principal components to account for genetic-ancestry admixture. Variant-Gene associations were deemed statistically significant if they reached a false discovery rate (FDR)-adjusted p-value ≤ 0.05. Analyses were restricted to biallelic variants (e.g., single-nucleotide polymorphisms), based on their higher genotyping accuracy and reliability relative to multiallelic variants. This increased precision enhances the robustness of our eQTL findings. Furthermore, the binary allele composition of biallelic variants facilitates interpretation of genetic effects and streamlines identification and characterization of associations between specific alleles and mRNA levels.

To evaluate the robustness and biological relevance of identified cis-eQTLs, we conducted replication analyses in tissues from the GTEx Project selected a priori based on their relevance to the pathophysiology of preeclampsia. Notably, because placental tissue, the central organ in preeclampsia pathogenesis, is not available in GTEx, we prioritized tissues that capture key maternal systems involved in disease development, including vascular, renal, and immune compartments. Vascular tissues, specifically the *Artery Aorta*, *Artery Tibial*, and *Artery Coronary*, were included to reflect the central role of endothelial dysfunction and impaired vascular remodeling in preeclampsia. These tissues provide a relevant proxy for systemic maternal vascular biology, a hallmark of the disorder. The *Kidney Cortex* was incorporated due to the clinical and mechanistic importance of renal involvement in preeclampsia, particularly in relation to proteinuria and glomerular endothelial injury. Genetic regulation of gene expression in this tissue may capture pathways contributing to renal manifestations of the disease. Finally, *Whole Blood* was included both to ensure consistency with the discovery dataset and to capture systemic immune and inflammatory processes known to contribute to preeclampsia pathogenesis.

This targeted tissue selection enables replication of eQTL signals across biologically relevant maternal systems implicated in preeclampsia while explicitly acknowledging the absence of placental tissue in GTEx. Replication in GTEx was defined as a variant–gene association demonstrating (i) a concordant direction of effect (beta) with the discovery analysis, (ii) an FDR-adjusted p-value ≤ 0.05 in the discovery data, and (iii) a nominal p-value ≤ 0.05 in any tissue in GTEx.

### Identification of Structural Variants in Linkage Disequilibrium with eQTLs

To investigate whether structural variants (SVs) may underlie or contribute to the regulatory effects observed at prioritized loci, we assessed linkage disequilibrium (LD) between identified cis-eQTL variants and SVs derived from the NeuroSeq cohort. Specifically, we leveraged high-confidence SV calls generated from long-read whole-genome sequencing of 30 individuals to capture genomic variation not represented in standard SNP-based GWAS datasets.

For each prioritized eQTL variant identified in the eQTL analysis, we extracted corresponding genotype information across the NeuroSeq samples. In parallel, genotype data for all detected SVs within the same individuals were compiled into a joint dataset. Merging SNPs and SVs into a single VCF is required because PLINK operates on unified variant sets to compute pairwise associations. SNPs were extracted based on genomic regions surrounding eQTL variants, and the merged VCF therefore contained both SNP and SV variants, enabling direct comparison of their linkage patterns.

LD between SNP-based eQTL variants and SVs was quantified using pairwise correlation metrics, specifically the squared correlation coefficient (r²), calculated across matched samples. Analyses were restricted to cis-regions, defined as ±1 Mb surrounding each eQTL variant, to prioritize local regulatory relationships. SVs demonstrating moderate-to-high LD with eQTL variants (r² ≥ 0.8) were considered putative candidates that may tag or functionally contribute to the observed gene expression associations. This approach enables the identification of SVs that may represent the true causal variants underlying GWAS and eQTL signals, particularly in genomic regions where SNPs serve only as proxies due to the limited resolution of short-read genotyping platforms.

Candidate SVs identified through LD analysis were further annotated using the NeuroSeq structural variant call set. For each SV, genomic coordinates, SV type, SV length, and allele frequency estimates were extracted from the jointly genotyped pbsv VCF generated from long-read whole-genome sequencing data. Structural variants were classified according to pbsv annotations, including insertions and deletions.

To evaluate the potential regulatory relevance of candidate SVs, genomic annotation analyses were performed using GENCODE gene models (release 44, GRCh38). Structural variants were intersected with annotated gene coordinates using BEDTools to identify SVs overlapping gene bodies. Promoter-overlap analyses were additionally performed using regions spanning ±2 kb from transcription start sites (TSS). For SVs not directly overlapping annotated genes or promoters, the nearest gene and genomic distance were determined using BEDTools closest.

Based on these annotations, SVs were classified into four genomic context categories: promoter-proximal, gene-body, proximal, and intergenic/distal. Promoter-proximal SVs were defined as variants overlapping promoter regions within ±2 kb of annotated TSS coordinates. Gene-body SVs represented variants directly overlapping annotated gene coordinates. Proximal SVs were defined as non-overlapping variants located within 10 kb of the nearest annotated gene, whereas intergenic/distal SVs represented variants located more than 10 kb from the nearest annotated gene.

By integrating long-read SV data with SNP-based regulatory associations, this framework provides a more comprehensive view of the genetic architecture and facilitates the discovery of previously unrecognized structural mechanisms that contribute to preeclampsia.

## RESULTS

### GWAS data

GWAS summary statistics were obtained from the GWAS Catalog for preeclampsia (EFO:0000668), essential hypertension (EFO:0000537), systolic blood pressure (SBP; EFO:0006335), and diastolic blood pressure (DBP; EFO:0006336). For each phenotype, we selected large-scale studies with publicly available data and sufficient genomic coverage and harmonized all variants to the GRCh38 reference genome to ensure consistent genomic positioning. Variants meeting a significance threshold of P ≤ 1 × 10⁻⁴ were extracted (**Table 1**). From these, we retained variants associated only with preeclampsia for subsequent analyses.

**Table 1:** Overview of GWAS datasets used in this study. For each phenotype (HTN, SBP, DBP, and PE), we report the total number of GWAS studies obtained from the GWAS Catalog along with the number of variants surpassing the predefined significance cutoff (P ≤ 1 × 10⁻⁴).

| Phenotype | Number of GWAS studies | Number of variants with p-value $\leq 0.0001$ |
| --- | --- | --- |
| Hypertension (HTN) | 117 | 663197 |
| Systolic Blood Pressure (SBP) | 112 | 1065509 |
| Diastolic Blood Pressure (DBP) | 83 | 841752 |
| Preeclampsia (PE) | 20 | 51750 |

### Preeclampsia-specific GWAS variants

After applying the significance threshold (P ≤ 1 × 10⁻⁴) to GWAS results, we focused specifically on variants uniquely associated with preeclampsia, rather than those shared across traits. From this set, we identified 10,843 preeclampsia-specific variants that were present in our whole-genome sequencing dataset. Variants with a minor allele frequency (MAF) ≥ 0.01 were retained and carried forward for subsequent eQTL discovery analyses.

### Demographic and clinical characteristics of study participants

We used whole-blood transcriptome and genotype data from 180 women for the eQTL analysis. **Table 2** summarizes participants’ demographic and clinical characteristics, including age, blood pressure, kidney function (estimated glomerular filtration rate, eGFR), anthropometric measures, lipid profile, and behaviors such as smoking and alcohol consumption.

**Table 2:** Demographic, clinical, and socio-behavioral characteristics of the study population (n = 180). Continuous variables are presented as mean ± standard deviation (SD), and categorical variables as counts and proportions.

| Characteristics | Mean/Count | SD/Proportion |
| --- | --- | --- |
| Age (years) | 48 | 12 |
| **HTN** |  |  |
| No | 22 | 12% |
| Yes | 158 | 88% |
| Systolic Blood Pressure (mm Hg) | 174 | 37 |
| Diastolic Blood Pressure (mm Hg) | 74 | 10 |
| Estimated Glomerular Filtration Rate (mL/min/1.73 m<sup>2</sup>) | 97 | 21 |
| Body Mass Index | 32 | 7 |
| Low Density Lipoprotein (mg/dL) | 114 | 44 |
| High Density Lipoprotein (mg/dL) | 62 | 14 |
| Triglycerides (mg/dL) | 76 | 38 |
| Glucose (mg/dL) | 102 | 31 |
| C-Reactive Protein (mg/L) | 5 | 8 |
| **Education Level** |  |  |
| ≤ High School | 55 | 31% |
| High School + Vocational Educational | 54 | 30% |
| ≥ Graduate Degree | 71 | 39% |
| **Current Smoker** |  |  |
| No | 170 | 94% |
| Yes | 10 | 6% |
| **Drinks Alcohol** |  |  |
| No | 67 | 37% |
| Yes | 113 | 63% |

The study population comprised 180 individuals with a mean age of 48 ± 12 years. Most participants were hypertensive (88%), with only 12% classified as non-hypertensive. Mean systolic and diastolic blood pressures were 174 ± 37 mm Hg and 74 ± 10 mm Hg, respectively. Kidney function was within the normal range, with a mean estimated glomerular filtration rate of 97 ± 21 mL/min/1.73 m². Participants were, on average, in the obese range (BMI: 32 ± 7).

Lipid, metabolic, and inflammatory markers showed variability across the cohort. Socio-behavioral characteristics indicated a diverse educational background, with the majority of participants being non-smokers and approximately two-thirds reporting alcohol consumption.

### eQTL results and validation in GTEx

The eQTL analysis was performed on expression profiles of 17,947 protein-coding genes, testing associations with 10,843 GWAS variants associated with Preeclampsia but not with Hypertension, SBP, and DBP, present in our whole-genome sequencing data with a MAF ≥ 0.01. Using a significance criterion of FDR-adjusted p-value ≤ 0.05, we identified 480 eQTL-mRNA associations, involving 277 unique variants and unique 192 protein-coding genes. The full list of eQTL-Gene pairs is reported in **Table S1** in the **Supplemental Material**.

To contextualize the regulatory signals identified from preeclampsia-specific GWAS variants, we examined their presence within the GTEx Project (version 8), a comprehensive resource for mapping cis-eQTLs across diverse human tissues. GTEx provides a robust framework for external validation, particularly as whole blood, the tissue used in our discovery analysis, is among the most well-powered datasets in the consortium, enabling reliable cross-study comparison.

For replication, we prioritized tissues with clear relevance to blood pressure regulation and vascular biology. Specifically, we included arterial tissues (aorta, tibial, and coronary) to capture mechanisms related to vascular tone and remodeling, kidney cortex to reflect renal control of blood pressure, and whole blood to enable direct validation of regulatory signals identified in our discovery dataset while also capturing systemic immune and inflammatory processes. Although placental tissue is highly relevant to the pathophysiology of preeclampsia, it is not currently represented in the GTEx resource, and therefore could not be included in our replication framework.

Replication analysis in GTEx supported 69 eQTL-gene associations derived from PE-specific variants, encompassing 35 variants and 14 genes across the five selected tissues (**Table S2** in the **Supplemental Material**). Notably, the majority of these signals (9 of 14 loci) demonstrated consistent regulatory effects across two or more tissues, particularly within vascular and renal contexts, underscoring the robustness and cross-tissue relevance of these PE-specific regulatory mechanisms (**Table 3**).

**Table 3:** Genes for which eQTL-gene pairs were identified, along with the number and types of tissues in which these regulatory associations were observed.

| Gene | Number of tissues where eQTL-Gene pair is observed | Tissue |
| --- | --- | --- |
| CPNE1 | 5 | Artery Aorta; Artery Coronary; Artery Tibial; Kidney Cortex; Whole Blood |
| NMRAL1 | 4 | Artery Aorta; Artery Coronary; Artery Tibial; Whole Blood |
| ZFP57 | 4 | Artery Aorta; Artery Coronary; Artery Tibial; Whole Blood |
| PLEKHA8 | 3 | Artery Aorta; Artery Tibial; Whole Blood |
| SULT1A1 | 3 | Artery Aorta; Artery Tibial; Whole Blood |
| TMEM116 | 3 | Artery Coronary; Artery Tibial; Whole Blood |
| JUP | 2 | Artery Aorta; Artery Tibial |
| TMEM204 | 2 | Artery Aorta; Whole Blood |
| TSGA10 | 2 | Artery Tibial; Whole Blood |
| ALG1L2 | 1 | Whole Blood |
| HLA-DPB1 | 1 | Artery Aorta |
| MICA | 1 | Artery Tibial |
| SEMA4D | 1 | Whole Blood |
| SULT1A2 | 1 | Whole Blood |

Across the five selected tissues, the distribution of significant eQTL-gene associations revealed a clear predominance in vascular and circulating compartments (Table 4). *Artery Tibial* showed the highest number of associations (n = 29), followed by *Whole Blood* (n = 24) and *Artery Aorta* (n = 10), with fewer signals observed in Artery Coronary (n = 5) and minimal representation in *Kidney Cortex* (n = 1). This pattern highlights a strong enrichment of preeclampsia-specific regulatory signals within vascular tissues, consistent with the central role of endothelial dysfunction and vascular remodeling in preeclampsia.

**Table 4:**
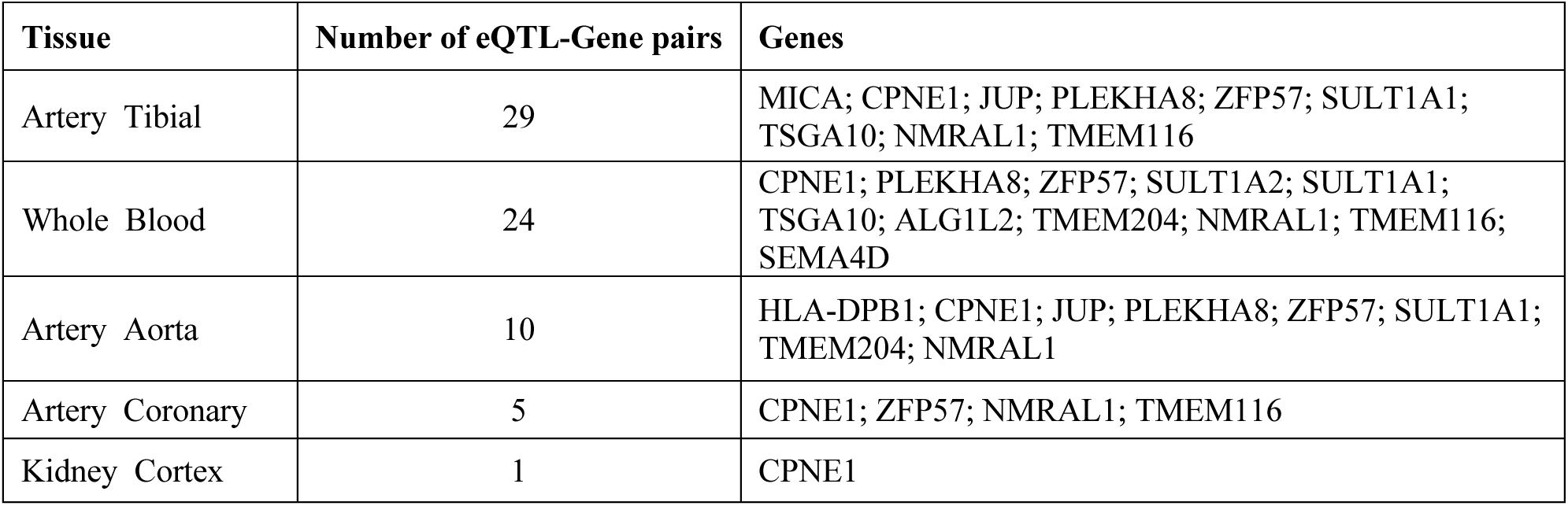
the number of significant eQTL-gene pairs identified in each tissue, ranked by total associations, along with the corresponding genes involved in these regulatory relationships.

| Tissue | Number of eQTL-Gene pairs | Genes |
| --- | --- | --- |
| Artery Tibial | 29 | MICA; CPNE1; JUP; PLEKHA8; ZFP57; SULT1A1; TSGA10; NMRAL1; TMEM116 |
| Whole Blood | 24 | CPNE1; PLEKHA8; ZFP57; SULT1A2; SULT1A1; TSGA10; ALG1L2; TMEM204; NMRAL1; TMEM116; SEMA4D |
| Artery Aorta | 10 | HLA-DPB1; CPNE1; JUP; PLEKHA8; ZFP57; SULT1A1; TMEM204; NMRAL1 |
| Artery Coronary | 5 | CPNE1; ZFP57; NMRAL1; TMEM116 |
| Kidney Cortex | 1 | CPNE1 |

Several genes were consistently identified across multiple tissues, including CPNE1, ZFP57, NMRAL1, PLEKHA8, and TMEM116, indicating shared regulatory mechanisms that are not tissue-restricted. In contrast, other signals appeared more context-specific, such as SEMA4D and ALG1L2 in whole blood and JUP in arterial tissues, suggesting complementary roles in immune, metabolic, and vascular pathways. The limited number of associations observed in kidney cortex suggests that, within the scope of preeclampsia-specific variants captured in this analysis, regulatory effects may be more prominently mediated through vascular and systemic pathways rather than renal-specific mechanisms.

### GWAS and eQTL effect integration

To integrate GWAS signals for preeclampsia with eQTL-derived regulatory effects, we performed a gene-level synthesis of the results, summarized in **Table 5**. Because multiple variants mapping to the same gene may be in linkage disequilibrium, effects were aggregated at the gene level rather than interpreted individually. GWAS effect directions were harmonized such that alleles associated with increased preeclampsia risk were consistently aligned, while the direction of eQTL effects reflects their impact on gene expression. No assumption was made regarding whether increased or decreased expression is inherently deleterious or protective; instead, the analysis evaluates whether preeclampsia risk alleles are associated with consistent regulatory shifts at each locus.

**Table 5:**
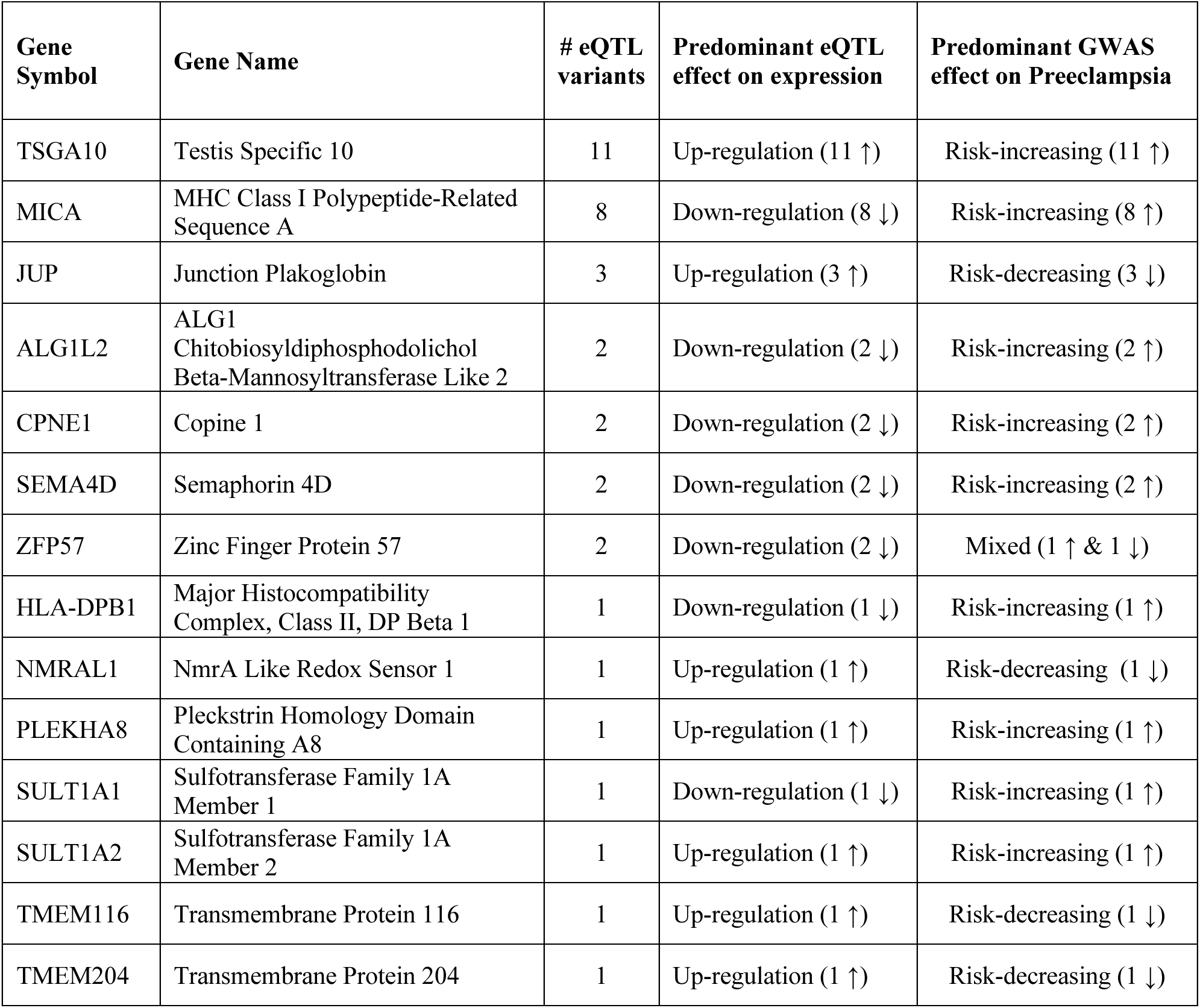
This table presents genes identified through cis-eQTL analysis of preeclampsia (PE)-specific GWAS variants, including the number of associated eQTLs per gene, the predominant direction of their effect on gene expression, and the corresponding direction of GWAS effects on disease risk. Arrows indicate the direction of effect (↑ increase; ↓ decrease), and “mixed” denotes inconsistent effects across variants.

This integration revealed several notable patterns. TSGA10 showed the largest number of eQTLs (n = 11), all associated with increased expression and concordant increases in preeclampsia risk. Similarly, JUP (n = 3) and PLEKHA8 (n = 1) displayed upregulation linked to risk-increasing alleles. In contrast, multiple genes, including MICA (n = 8), ALG1L2, CPNE1, and SEMA4D (each n = 2), exhibited predominant downregulation despite being associated with increased preeclampsia risk, indicating inverse relationships between expression and genetic risk at these loci.

Most genes with fewer eQTLs (n = 1) also showed consistent patterns of downregulation paired with risk-increasing alleles, including MICA, HLA-DPB1, and SULT1A1, although some signals (e.g., NMRAL1, TMEM116, and TMEM204) suggested discordant or less consistent directionality between expression and GWAS effects. Notably, ZFP57 was the only gene demonstrating mixed GWAS effects despite consistent downregulation, reflecting heterogeneity in the direction of genetic associations across variants. Full details of all variant-gene-trait relationships are provided in **Table S3** in the **Supplemental Material**.

### Structural variants in Linkage Disequilibrium with eQTL loci

To further investigate potential structural mechanisms underlying replicated regulatory associations, LD analyses were repeated using the subset of replicated GTEx-supported eQTL variants. This analysis identified 66 high-LD SNP-SV-gene associations involving 12 replicated eQTL variants, 8 candidate structural variants, and 3 replicated eGenes represented within the final SV-linked loci. Most identified associations demonstrated strong to perfect LD (r² ≥ 0.8), including several perfect proxy relationships (r² = 1), suggesting that these SVs may represent candidate causal variants underlying observed SNP-based regulatory signals.

Structural variant annotation revealed that the candidate loci consisted of both insertions and deletions identified from long-read sequencing data generated in the NeuroSeq cohort. Insertions represented the majority of identified associations, accounting for 46 associations involving 3 unique SVs, while deletions accounted for 20 associations involving 5 unique SVs.

Genomic context annotation demonstrated that most candidate SVs localized within regulatory or gene-associated regions. Specifically, gene-body overlaps represented the largest category, accounting for 38 associations involving 5 unique SVs, 12 unique eQTL variants, and 3 eGenes. Promoter-proximal SVs accounted for 24 associations involving 2 unique SVs and 6 unique eQTL variants, while proximal SVs located near annotated genes accounted for 4 associations involving 1 unique SV and 1 unique eQTL variant. No candidate SVs were classified as distal/intergenic, indicating that all prioritized loci localized within or near annotated gene regulatory regions.

Several candidate SV loci clustered within biologically relevant genomic regions. A prominent cluster of insertion events on chromosome 2 demonstrated strong LD with replicated eQTL variants associated with TSGA10 regulatory signals across both whole blood and arterial tissues. Additional candidate loci were identified within the chromosome 6 major histocompatibility complex (MHC) region, including structural variants linked to immune-related regulatory loci. Multiple deletion events on chromosome 16 demonstrated strong LD with replicated eQTL variants regulating SULT1A1 and SULT1A2 expression in vascular and whole blood tissues.

Together, these findings demonstrate that replicated regulatory variants associated with preeclampsia-related loci frequently localize to regions harboring structurally complex genomic variation, supporting the hypothesis that structural variants may contribute to the molecular mechanisms underlying observed GWAS and eQTL signals.

Representative candidate SV loci, including SV type, allele frequency, genomic context, nearest genes, and linked replicated eQTL signals, are summarized in **Table 6**. Full details of all SNP-SV-gene associations identified through the LD analysis are provided in **Table S4** in the **Supplemental Material**.

**Table 6:** Structural variants identified from long-read sequencing data that are in high linkage disequilibrium with replicated eQTL loci associated with candidate genes. The table summarizes structural variants (SVs) detected in the NeuroSeq long-read sequencing dataset that showed high linkage disequilibrium (LD; r. ^2^ ≥ 0.80) with replicated eQTL variants. For each SV, the associated gene(s), GTEx tissues in which the eQTL signal replicated, number of linked eQTL variants, LD strength, SV class, SV size, allele frequency in the NeuroSeq cohort, and genomic context are reported.

| Structural Variant (SV) ID | Gene associated (via eQTL) | GTEx Tissue Replicated | Number of eQTL in LD with SV | LD ($r^2$) | SV Type | SV size (bp) | SV Frequency in NeuroSeq | SV Genomic Context |
| --- | --- | --- | --- | --- | --- | --- | --- | --- |
| chr2_99279416_INS | TSGA10 | Whole Blood;<br>Artery Tibial | 11 | 1 | INS | 40 | 0.35 | gene-body |
| chr2_99338993_INS | TSGA10 | Whole Blood;<br>Artery Tibial | 6 | 0.94 | INS | 22 | 0.33 | promoter-proximal |
| chr2_99490085_INS | TSGA10 | Whole Blood;<br>Artery Tibial | 6 | 0.94 | INS | 22 | 0.67 | promoter-proximal |
| chr16_28380189_DEL | SULT1A2;<br>SULT1A1 | Whole Blood;<br>Artery Aorta;<br>Artery Tibial | 1 | 0.86 | DEL | 51 | 0.19 | gene-body |
| chr16_28448926_DEL | SULT1A2;<br>SULT1A1 | Whole Blood;<br>Artery Aorta;<br>Artery Tibial | 1 | 0.85 | DEL | 86 | 0.17 | proximal |
| chr16_28698579_DEL | SULT1A2;<br>SULT1A1 | Whole Blood;<br>Artery Aorta;<br>Artery Tibial | 1 | 0.84 | DEL | 40 | 0.29 | gene-body |
| chr16_28465698_DEL | SULT1A2;<br>SULT1A1 | Whole Blood;<br>Artery Aorta;<br>Artery Tibial | 1 | 0.84 | DEL | 169 | 0.14 | gene-body |
| chr16_28351857_DEL | SULT1A2;<br>SULT1A1 | Whole Blood;<br>Artery Aorta;<br>Artery Tibial | 1 | 0.80 | DEL | 169 | 0.15 | gene-body |

## DISCUSSION

Preeclampsia (PE) sits at the intersection of placental, immune, and vascular pathophysiology, and decades of genetic research have produced an architecture that substantially overlaps with that of essential hypertension and quantitative blood pressure traits^14, 29^. When shared signals dominate, however, mechanisms uniquely engaged by PE, particularly those tied to placentation and maternal-fetal immune adaptation, can be obscured. By restricting our discovery set to variants associated with PE but not with hypertension, systolic, or diastolic blood pressure, we sought to enrich for regulatory signals more proximally linked to disease-specific biology. The resulting picture is consistent with a model in which PE-specific susceptibility converges on maladaptive maternal-fetal immune crosstalk and systemic endothelial regulation, with structural variation contributing to the regulatory architecture at several loci.

### MHC signals implicate maladaptive maternal-fetal immune signaling

The strongest replicated PE-specific signals localized to the major histocompatibility complex (MHC) on chromosome 6, a region with a long-established role in reproductive immunology^35, 41–43^. The downregulation of *MICA* in arterial tissue among PE risk allele carriers is mechanistically coherent with decidual natural killer (dNK) cell biology: MICA is a ligand for the activating receptor NKG2D expressed on dNK cells, and the dNK-trophoblast interaction is a determinant of spiral artery remodeling in early pregnancy^44–47^. Reduced *MICA* expression on PE risk alleles plausibly attenuates this signal, impairing the controlled cytotoxicity and cytokine output that drive remodeling and producing the placental ischemic phenotype that initiates downstream maternal disease.

The *HLA-DPB1* signal in artery aorta extends the immune axis from innate to adaptive compartments. Pregnancy requires tightly orchestrated tolerance of fetal antigens^48–50^, and HLA class II variation has been implicated repeatedly in PE susceptibility^41–43^. That this eQTL replicated in arterial rather than placental or immune tissue suggests that MHC-mediated effects on maternal vascular biology may extend systemically beyond the placental compartment.

The *SEMA4D* signal, replicated in whole blood, links these immune findings to vascular biology. Semaphorin 4D is now established as a mediator of endothelial-macrophage crosstalk, including macrophage polarization and angiogenic responses^51^. Given the central role of angiogenic imbalance in PE^52^, dysregulation of *SEMA4D* may represent one molecular bridge between altered immune signaling and impaired vascular adaptation. Together, these findings support the contemporary view that PE is a disorder of disrupted immunovascular communication at the maternal-fetal interface, with effects extending beyond the placenta into systemic maternal vasculature^13, 48, 53^.

### Vascular tissue enrichment supports systemic endothelial involvement

Replicated eQTL signals were enriched in artery tibial (n = 29 pairs) and whole blood (n = 24) relative to kidney cortex (n = 1). While GTEx kidney cortex is less well-powered than the arterial panels, the magnitude of the imbalance, together with multi-tissue replication in 9 of 14 prioritized loci, argues against a pure power artefact. PE-specific regulatory effects appear to operate substantially through systemic maternal endothelium, consistent with the conceptualization of PE as a generalized endothelial disorder driven by circulating anti-angiogenic factors^54^, in which renal manifestations reflect glomerular endothelial injury downstream rather than primary renal dysfunction.

Two vascular hits warrant specific comment. *JUP* (junction plakoglobin) encodes a core component of adherens junctions and desmosomes that contributes to endothelial barrier integrity and cell-cell adhesion^55^; the upregulation observed on PE risk alleles may perturb Wnt signaling and adherens-junction composition, with downstream effects on endothelial mechanosensing. *TMEM204* has been associated with endothelial biology and tumor vascular phenotypes^56^ but is essentially uncharacterized in pregnancy, making it a candidate worth pursuing in trophoblast-endothelial models.

### An imprinting-maintenance signal extends PE genetics into placental epigenetics

The replication of regulatory signals at *ZFP57* is, in our view, one of the more conceptually novel findings of this work. *ZFP57* is the master maintainer of genomic imprints during early development^57, 58^, and the placenta is unusually dependent on imprinted gene expression: imprinted clusters at 11p15.5 (*IGF2/H19*), 14q32, and elsewhere are critical determinants of trophoblast invasion, placental size, and fetal growth^59^. Disruption of imprinting maintenance has been linked experimentally to placental insufficiency phenotypes that overlap mechanistically with PE^60^. The identification of replicated PE-specific eQTLs at *ZFP57* therefore positions imprinting alongside immunity and vascular biology as a third mechanistic axis and suggests a testable hypothesis: that allelic variation at *ZFP57* perturbs imprint maintenance at placentally relevant loci, with downstream consequences for trophoblast function.

### Structural variants nominate candidate causal alleles at SNP-tagged loci

A central methodological contribution of this work is the integration of long-read structural variants (SVs) with SNP-based eQTL signals. The 66 high-LD (r² ≥ 0.80) SNP-SV-gene relationships identified here suggest that several replicated regulatory signals may tag structural rather than point variation^31–33^. The chromosome 2 insertion cluster in strong LD with *TSGA10* eQTLs is illustrative: the lead SV (chr2_99279416_INS, r² = 1) localizes to the gene body, with two additional candidate insertions within the ±2 kb promoter window of the *TSGA10* transcription start site, a configuration consistent with cis-regulatory disruption in a region known to harbor complex structural variation poorly resolved by short-read sequencing^30, 31^.

The chromosome 16 deletion cluster linked to *SULT1A1*/*SULT1A2* is mechanistically attractive for an independent reason: the 16p11.2 region contains well-characterized segmental duplications that produce extensive copy-number variation across sulfotransferase paralogs, and altered estrogen sulfation has been proposed as a contributor to PE pathophysiology. Identification of multiple deletions in high LD with *SULT1A1*/*SULT1A2* eQTLs is consistent with copy-number variation underlying the observed expression effects, generating a hypothesis that could not have been derived from short-read SNP data alone. More broadly, these findings illustrate how long-read sequencing in African-ancestry populations, which harbor the greatest structural genomic diversity^34^, can recover regulatory architecture systematically under-ascertained by conventional GWAS pipelines^31–33^.

### Strengths and limitations

The principal strengths of this work are the integration of GWAS, transcriptomic, cross-tissue eQTL replication, and long-read SV discovery within a single analytical framework, and the focus on an African-ancestry population disproportionately affected by PE yet historically underrepresented in genomic research^26–28, 34^. Several limitations should be addressed. First, the discovery transcriptome cohort comprised non-pregnant, predominantly hypertensive women; whole-blood eQTLs serve as informative proxies for systemic regulation but cannot directly capture placenta- or pregnancy-specific programs, and the hypertensive state may modulate expression beyond what age and ancestry adjustments capture. Second, the long-read SV cohort (n = 30) is small. and ancestry-heterogeneous. Although this ancestral mismatch may influence local LD structure at some loci, the use of an African-ancestry long-read reference panel remains appropriate for identifying common candidate SVs linked to regulatory variants in African-ancestry genomes. Third, the construct of “PE-specific” should be read as operational rather than absolute: because the PE GWAS corpus is smaller and less well-powered than that for blood pressure traits, our filter preferentially retains variants where blood pressure GWAS lacked power to detect shared signals, and some prioritized loci may, in larger analyses, prove shared with broader cardiovascular phenotypes. Finally, functional validation in trophoblast, endothelial, and immune-cell models will be required to establish causality.

## Conclusions

The findings support a model in which PE-specific genetic susceptibility converges on three interlinked mechanistic axes: MHC-anchored maternal-fetal immune signaling, systemic maternal vascular regulation, and imprinting-related epigenetic maintenance, with long-read SV data nominating candidate causal alleles missed by conventional GWAS pipelines. Priorities for follow-up include replication of the eQTL signals in pregnancy-specific tissues (placenta, decidua, and trophoblast), larger long-read studies in African-ancestry pregnancy cohorts, and functional dissection of the prioritized loci, particularly *MICA*, *HLA-DPB1*, *ZFP57*, and the *SULT1A* copy-number locus. More broadly, the work illustrates the analytical gains from combining trait-specificity-aware variant prioritization, multi-tissue regulatory mapping, and structural variant discovery in the diverse populations that bear the heaviest disease burden.

## ACKNOWLEDGEMENTS

This work was supported by the Chan Zuckerberg Initiative’s Foundation to Accelerate Precision Health Program and Advance Genomics Research at Meharry Medical College (CZIF2022-007043) and by the NIMHD grant U54MD007593 to build research capacity at Meharry Medical College.

We thank Prof Jeannine Heckmann, Prof Andre Mochan, Prof Dilraj Sokhi, Prof Adesola Ogunniyi and their clinical teams at ALSAfrica Network sites (https://alsafrica.org) for their contributions to participant recruitment and sample collection for the NeuroSeq study.

We acknowledge the use of the ilifu cloud computing facility (www.ilifu.ac.za), a partnership between the University of Cape Town, the University of the Western Cape, Stellenbosch University, Sol Plaatje University, and the Cape Peninsula University of Technology.

